# Glycine homeostasis requires reverse SHMT flux

**DOI:** 10.1101/2023.01.11.523668

**Authors:** Matthew J. McBride, Craig J. Hunter, Joshua D. Rabinowitz

## Abstract

**SUMMARY:** The folate-dependent enzyme serine hydroxymethyltransferase (SHMT) reversibly converts serine into glycine and a tetrahydrofolate-bound one-carbon unit. Such one-carbon unit production plays a critical role in development, the immune system, and cancer. Here we show that the whole-body SHMT flux acts to net consume rather than produce glycine. Pharmacological inhibition of whole-body SHMT1/2 and genetic knockout of liver SHMT2 elevated circulating glycine levels up to eight-fold. Stable isotope tracing revealed that the liver converts glycine to serine, which is then converted by serine dehydratase into pyruvate and burned in the tricarboxylic acid cycle. In response to diets deficient in serine and glycine, de novo biosynthetic flux was unaltered but SHMT2- and serine dehydratase-mediated catabolic flux was lower. Thus, glucose-derived serine synthesis does not respond to systemic demand. Instead, circulating serine and glycine homeostasis is maintained through variable consumption, with liver SHMT2 as a major glycine-consuming enzyme.

## INTRODUCTION

The non-essential amino acids serine and glycine are circulating nutrients required by mammalian tissues to support biosynthesis of proteins, nucleotides, and lipids^1^. Homeostasis of these amino acids requires balancing production (by dietary intake and de novo synthesis) with consumption (by biosynthetic and catabolic pathways)^2^. Depletion of circulating glycine and thus disrupted homeostasis is a hallmark of metabolic syndrome^3,4^.

De novo synthesis of both serine and glycine begins from the glycolytic intermediate 3-phosphoglycerate via three enzymatic reactions initiated by phosphoglycerate dehydrogenase (PHGDH) to make serin^5–7^. Serine is then transformed by serine hydroxymethyltransferase in the cytoplasm (SHMT1) or mitochondria (SHMT2) to glycine and a one-carbon (1C) unit bound to tetrahydrofolate (THF)^8^. Glycine can also be synthesized from threonine catabolism, though this pathway is lost in humans^9^. In cell lines, depletion of serine and glycine from culture media drives an increase in their de novo synthesis from glucose^10,11^. This is consistent with intracellular de novo synthesis of non-essential amino acids increasing when available supply from the extracellular environment is reduced.

A major function of serine and glycine is to provide 1C units for purine, thymidine, and methionine synthesis^12–14^. This can occur through serine conversion to glycine plus a 1C unit by SHMT or through glycine conversion to CO_2_ plus a 1C unit by glycine cleavage system (GCS)^15^. Among the two SHMT isozymes, SHMT2 is more functionally important. Genetic deletion of either SHMT2 or GCS results in development defects prototypical of folate or 1C deficiency^16,17^. Outside of development, SHMT2 is a key driver of T cell activation and cancer cell proliferation^18,19^.

While most interest in serine and glycine metabolism has focused on their importance as biosynthetic precursors, there is also need to catabolize these amino acids to prevent excessive accumulation. Classically, serine is catabolized to pyruvate by the liver-specific enzyme serine dehydratase^20^ and to glycine by SHMT1/2 with glycine subsequently cleared by GCS in the liver^21,22^. Humans with a biallelic loss-of-function mutation in GCS develop non-ketotic hyperglycinemia (NKH) characterized by plasma glycine concentrations up to nine times greater than normal^23^. A third of these patients do not to survive a full year and glycine level correlates with disease severity. Those patients that do reach infancy have neurological impairments that include seizures and diminished psychomotor development.

While SHMT2 is reversible and so can consume glycine and a 1C unit to produce serine, evidence to date suggested that all functional flux was in the forward (i.e. serine-consuming) direction. Surprisingly, however, when we tested pharmacological dual SHMT1/2 inhibitors in mice, rather than seeing glycine depletion as expected, we observed massive elevations. Combining mouse genetics, metabolomics, and tracing, we show that a major path of systemic glycine consumption is liver conversion to serine via SHMT2. This serine can then be further converted by serine dehydratase to pyruvate which can then be oxidized in the TCA cycle. We additionally explore how systemic serine and glycine homeostasis is achieved. Our data support a model of homeostasis where de novo synthesis is independent of dietary supply. Instead, consumption rates, including via the liver SHMT2 pathway, change in proportion to circulating levels.

## RESULTS

### Loss of hepatic SHMT2 elevates serum glycine

Classically, serine is synthesized from the glycolytic intermediate 3-phosphoglycerate via the PHGDH pathway, and catabolized by the enzyme serine dehydratase, whose expression is restricted primarily to liver (Figure S1A). Glycine is synthesized from serine via the folate-dependent enzyme serine hydroxymethyltransferase, and catabolized by another folate-dependent enzyme, the glycine cleavage system (Figure 1A, Figure S1B). We treated mice with vehicle or a dual SHMT1/2 small molecule inhibitor (SHIN2) by intravenous infusion for 12 hours^24^. Serum collected following treatment was analyzed by liquid chromatography-mass spectrometry (LC-MS). Rather than suppressing glycine levels, SHIN2 treatment increased circulating glycine from ~400 μM to over 3 mM (Figure 1B, Figure S1C). The other strongest metabolite changes were glycine-containing metabolites, such as acetylglycine and longer-chain acylglycines (Figure 1B). Thus, rather than being a net source of circulating glycine, SHMT1/2 appears to be a major glycine consumer.

**Figure 1:**
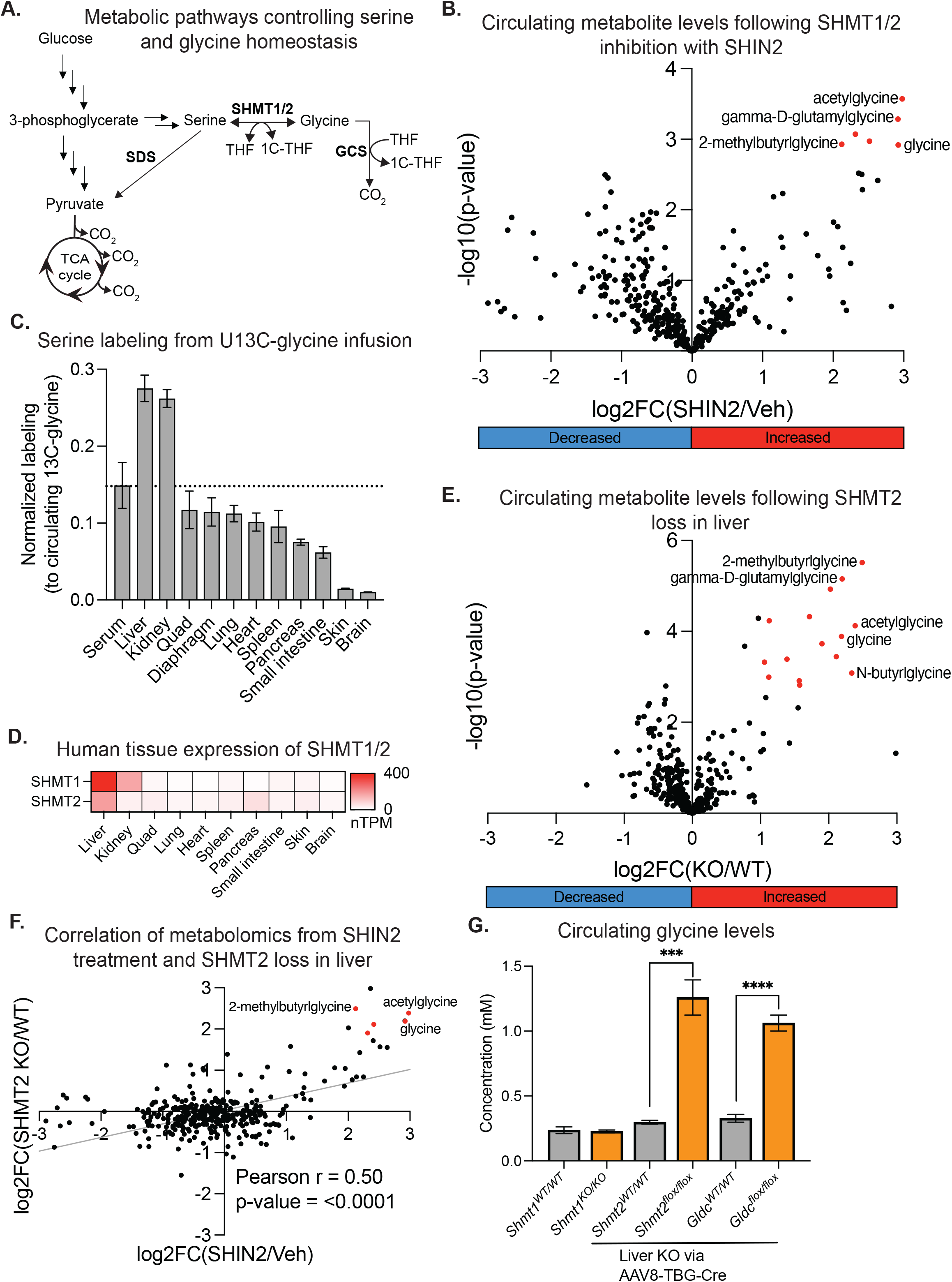
Hepatic SHMT2 is required for glycine clearance. (A) Serine and glycine production and consumption pathways. (B) Serum metabolomics from C57BL/6 mice treated with vehicle or SHIN2 (n=3). (C) Normalized serine labeling from U13C-glycine infusion in circulation and tissues for fed mice. (D) Gene expression level of SHMT1 and SHMT2 across tissues in humans. (E) Serum metabolomics from C57BL/6-Shmt2^flox/flox^ mice upon inducing SHMT2 liver knockout by AAV8-TBG-Cre viral infection (n=5-7). (F) Pearson correlation analysis of serum metabolomics from (B) and (E). (G) Serum glycine concentrations from C57BL/6 mice with whole-body SHMT1, liver-induced SHMT2 or liver-induced GLDC gene knockout (n=3-7). All data are reported as mean ± SEM, p value by unpaired T test and replicates (n) indicated. Metabolomics data underwent Benjamini-Hochberg correction for multiple comparison testing.

As small molecules can have off target effects, we turned to genetics to validate this finding and identify the isozyme and tissue responsible for this SHMT-mediated glycine consumption. Whole-body SHMT1 knockout has only modest phenotypes with no reported changes in circulating glycine levels, while whole-body SHMT2 knockout is embryonic lethal in mice^16,25,26^. Thus, we prioritized SHMT2, and generated a SHMT2-floxed mouse (Shmt2^flox/flox^ C57BL/6) for this purpose^27^.

To explore the most relevant tissue(s) in which to knockout SHMT2, we infused U13C-glycine systemically in mice and measured serine labeling in different tissues and in the circulation. We hypothesized that serine labeling from glycine would be high in the tissue(s) doing the bulk of reverse SHMT flux. Consistent with this, we observed high serine labeling from the infused U13C-glycine in both liver and kidney, while labeling in all other tissues was lower than in the circulation (Figure 1C). Thus liver and/or kidney are responsible for systemic glycine to serine flux. As both SHMT2 and the major serine catabolic enzyme serine dehydratase (SDS) are most highly expressed in liver (Figure 1D, Figure S1A), we prioritized liver. Using AAV8 viral infection, we induced liver-specific Cre expression to knockout SHMT2 in Shmt2^flox/flox^ C57BL/6 mice and collected serum for metabolomics. Similar to SHIN2 treatment, liver-specific SHMT2 knockout dramatically elevated serum glycine and related metabolites (Figure 1E-G).

As expected, whole-body SHMT1 knockout had no effect on circulating glycine levels (Figure 1G). Knockout of the canonical glycine-catabolic enzyme, glycine decarboxylase (GLDC) of the glycine cleavage system, in a liver-specific manner resulted in similar circulating glycine elevation to liver-specific SHMT2 loss^28^ (Figure 1G). Thus, glycine homeostasis requires both the glycine cleavage system and reverse flux through liver SHMT2.

### Glycine is converted to serine for hepatic clearance by TCA cycle oxidation

Reverse SHMT2 flux consumes glycine and a 1C unit to generate serine. In the liver, serine dehydratase can convert serine to pyruvate for consumption by TCA cycle oxidation (Figure 2A, Figure S1A). To determine the fate of circulating glycine, we infused U13C-glycine in rats and collected arterial and site-specific venous serum samples, which were used to measure metabolite concentrations and labeling by LC-MS (Figure 2B). In rat, the liver net consumed both serine and glycine, as evidenced by lower concentrations in the hepatic vein than incoming blood (22% artery & 78% portal vein) (Figure S2A)^29^. Moreover, the liver catabolized glycine to pyruvate to lactate, as evidenced by increased M+2 lactate labeling in the hepatic vein from U13C-glycine infusion (Figure 2C).

**Figure 2:**
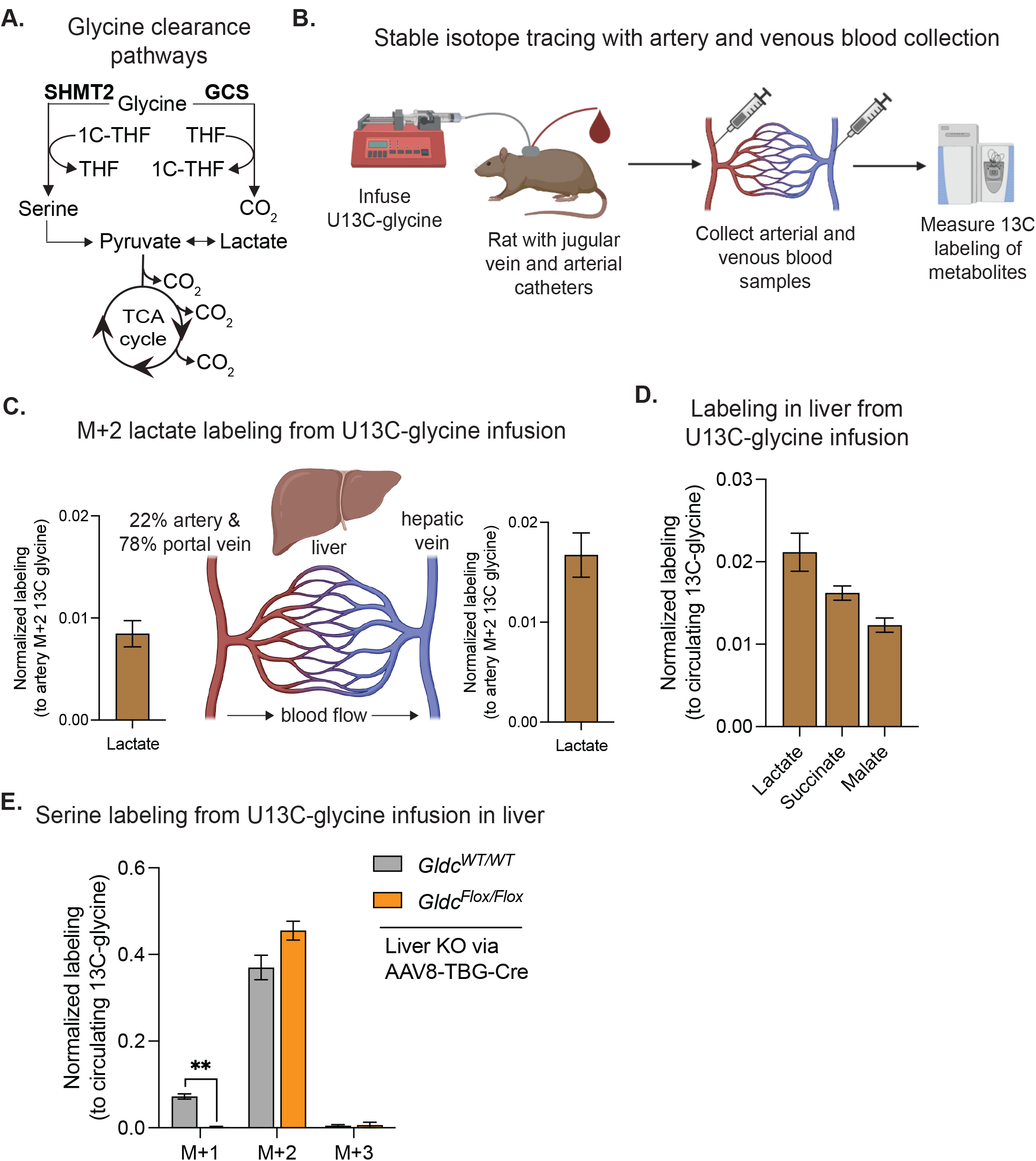
Reverse SHMT2 flux clears glycine by TCA oxidation in liver. (A) Pathways of glycine consumption. (B) Experimental diagram for artery and venous sampling following stable-isotope tracing in rats. (C) Artery and venous lactate labeling from U13C-glycine infusion in Sprague Dawley rats (n=4). (D) Normalized labeling from U13C-glycine infusion of lactate, succinate and malate in liver for fed mice. (E) Normalized serine labeling from U13C-glycine infusion in liver of wild-type and liver GLDC knockout mice (n=3). All data are reported as mean ± SEM, p value by unpaired T test and replicates (n) indicated.

To further explore the fate of glycine in the liver, we measured labeling from infused U13C-glycine into liver lactate and the TCA cycle intermediates (succinate and malate) (Figure 2D). This indicates that serine produced by reverse SHMT flux feeds via serine dehydratase into the TCA cycle. The hepatic labeling of serine, lactate and TCA cycle intermediates from glycine is lost with SHIN2 treatment (Figure S2B).

Reverse SHMT2 flux requires a 1C unit in the form of methylene-THF, and one source of methylene-THF is the glycine cleavage system. We infused U13C-glycine in both wildtype and liver GLDC knockout mice and measured liver serine labeling. Odd isotope labeling of serine (M+1 & M+3) occurs when unlabeled or M+2 labeled glycine condenses with M+1 labeled 1C unit made from glycine by the glycine cleavage system. Such odd-labeled serine was readily observable in wildtype mice and accounted quantitatively (after correction for the extent of glycine labeling) for 17% of liver 1C units (Figure 2E, Figure S2C). As expected, such labeling was lost with GLDC knockout. Loss of the glycine cleavage system, however, did not alter serine M+2 labeling from infused U13C-glycine, reflecting the sufficiency of other sources of methylene-THF to drive reverse SHMT2 flux. This is evidence that glycine clearance by reverse SHMT2 flux can occur independent of glycine cleavage system activity.

### Glycine- and serine-deficient diets decrease circulating levels selectively in fed state

In mammals, circulating serine and glycine comes from three sources: (i) dietary protein catabolism, (ii) endogenous protein breakdown, or (iii) de novo serine synthesis from the glycolytic intermediate 3-phosphoglycerate (Figure 3A). Serine- and glycine-free diets are well tolerated by mice for months and slow growth of some cancers^11,30,31^. Having uncovered an unexpected glycine homeostatic pathway (liver SHMT2 pathway), we turned to how serine and glycine homeostasis is maintained (Figure 3B). It has previously been reported that serine-and glycine-free diets decrease circulating levels of both amino acids by about 50%, which we also observed to be true in the fed state (Figure 3C). Interestingly, however, we observed no change in fasted mice (Figure 3C). Consistent with circulating levels, tissue pools were depleted in fed state and maintained in fasted state when mice consumed the ser/gly-free chow (Figure S3A-B). The minimalist interpretation is that the serine-and glycine-free diet is hardly altering endogenous metabolism, merely decreasing the dietary influx in the fed state. Indeed, we observed similar circulatory turnover fluxes (endogenous production rates) for both serine (no change) and glycine (modest decrease) across fasted mice on regular or serine-and glycine-free diet (Figure 3D).

**Figure 3:**
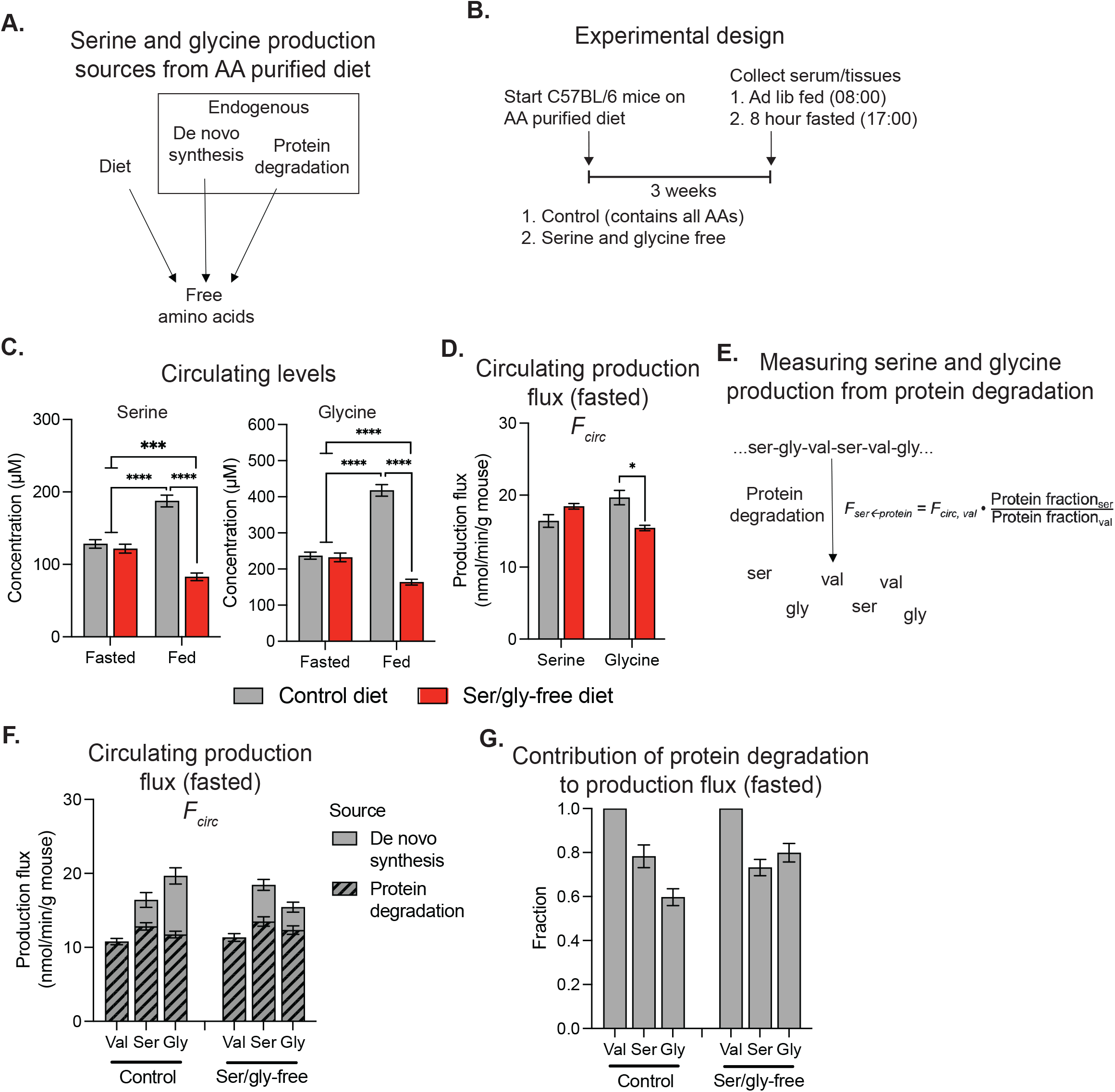
Diet-induced changes in circulating serine and glycine levels are lost in the fasted state when protein degradation is the major production source. (A) Schematic of free amino acid sources when mice consume a purified diet. (B) Experimental design. (C) Serum serine and glycine concentrations (n=13-17). (D) Serine and glycine turnover fluxes for fasted mice on control or ser/gly-free diet (n=4-8). (E) Method for quantifying serine or glycine production from protein degradation based on production flux of an essential amino acid (valine). (F) Valine, serine, and glycine turnover fluxes for fasted mice on control or ser/gly-free diet (n=4-10). (G) Fraction of circulating valine, serine and glycine production from protein degradation. All data are reported as mean ± SEM, p value by unpaired T test and replicates (n) indicated.

To explore the flux from endogenous protein catabolism to serine and glycine, we infused fasted mice with U13C-valine, U13C-glycine or U13C-serine to calculate turnover fluxes (*F_circ_*) for each amino acid. Because valine is an essential amino acid, its fasted production is from endogenous protein catabolism:

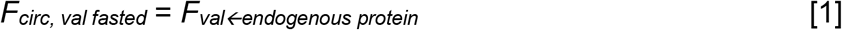

Protein catabolism generates all amino acids in proportion to their abundance in protein. For serine, this implies

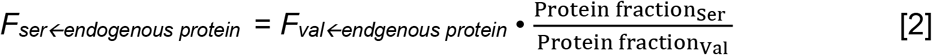

and similarly for glycine. By comparing these production fluxes from endogenous protein catabolism to overall circulatory flux, we found that, in the fasted state, 60-80% of both serine and glycine production is from protein degradation on both control and ser/gly-free diet (Figure 3F-G). Consequently, protein degradation rather than de novo synthesis is the major source of whole-body production of these amino acids in fasted mice.

### Loss of dietary source has no effect on endogenous production

Serine- and glycine-free diet selectively alters circulating serine and glycine levels in the fed state (Figure 3C). Consistent with this, we observed about 25% higher fed-state fluxes for both amino acids in mice receiving a full assortment of dietary amino acids, as opposed to a serine-and glycine-free diet (Figure 4A). We calculated flux from incoming dietary amino acids based on quantity of free serine and glycine contained in chow and total quantity of chow each mouse consumed per day (Figure S4A-B). Strikingly, after subtracting out incoming flux from diet, the remaining flux was unaltered on serine-and glycine-free diet (Figure 4B). This further supports endogenous metabolism being largely unaltered by dietary serine and glycine removal.

**Figure 4:**
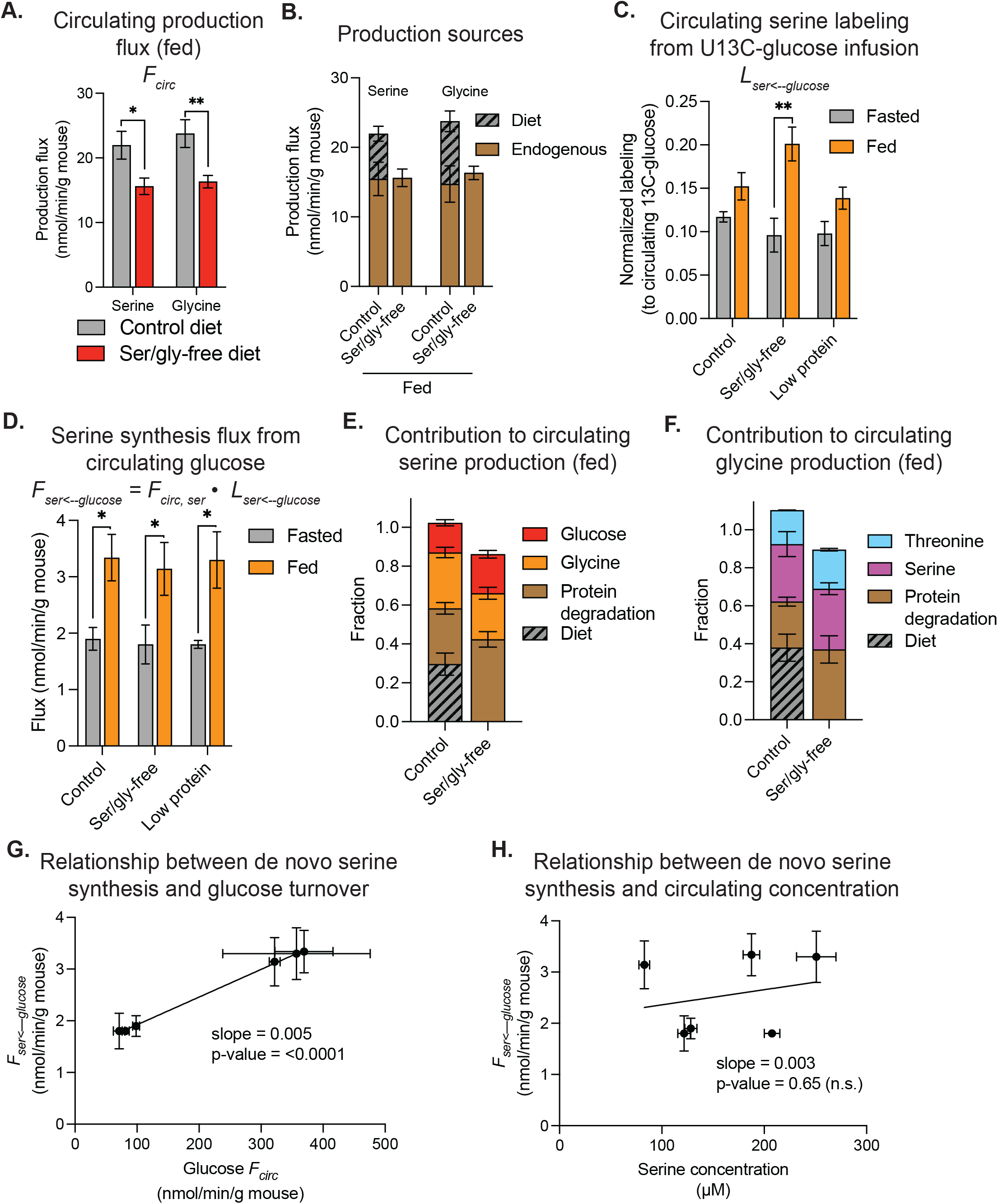
Feeding increases de novo serine synthesis from circulating glucose independent of serine levels. (A) Serine and glycine turnover fluxes for fed mice on control or ser/gly-free diet (n=7-9). (B) Proportion of serine and glycine production fluxes from diet and endogenous sources for fed mice on control or ser/gly-free diet. Incoming flux from diet was subtracted from total turnover flux (*F_circ_*) to determine endogenous fux. (C) Normalized serine labeling in circulation from U13C-glucose infusion. (D) De novo serine synthesis flux from circulating glucose. (E, F) Fraction of circulating (E) serine and (F) glycine production from each source for fed mice on control or ser/gly-free diet. (G) Relationship between de novo serine synthesis flux and glucose turnover flux. (H) Lack of relationship between de novo serine synthesis flux and circulating serine concentration. All data are reported as mean ± SEM, p value by unpaired T test and replicates (n) indicated.

To probe more explicitly de novo serine synthesis, we used U13C-glucose tracing to measure contribution to total serine production flux. The fractional fed-state labeling of circulating serine from U13C-glucose tracer was increased in mice receiving serine-and glycine-free diet (Figure 4C). This increase, however, could potentially simply reflect less dilution from dietary serine, as opposed to higher de novo synthesis. Indeed, multiplying the fraction of circulating serine coming from glucose by the serine circulatory turnover flux, to quantify the de novo synthetic flux, revealed no change between normal and serine-and glycine-free diet (Figure 4D, Figure S4C). Overall, the main sources of serine and glycine were diet, endogenous protein degradation, interconversion between these two amino acids or de novo synthesis of serine from glucose and glycine from threonine (Figure 4E-F). Notably, there does not appear to be mechanisms to upregulate serine synthesis in response to its dietary depletion.

We wondered if this might reflect the lack of specific mechanisms for sensing dietary serine, with physiological sensing instead focused on overall dietary protein. Feeding a low-protein diet (5%), however, similarly did not alter de novo serine synthetic flux (Figure 4C-D, Figure S4C). Overall, across the three dietary conditions in both fed and fasted states, we observed a trend toward higher serine synthesis from circulating U13C-glucose in the fed state, resulting in a correlation between circulating glucose flux and serine biosynthetic flux (Figure 4G). This may reflect in part higher 3-phosphoglycerate labeling from circulating glucose in fed mice. Most importantly, we observed no relationship between circulating serine levels and de novo synthesis flux from glucose (Figure 4H), implying the lack of a homeostatic mechanism to turn on endogenous serine synthesis when circulating serine is depleted.

### Homeostasis is maintained by modulating serine and glycine consumption

Since overall circulatory turnover fluxes of glycine and serine are decreased on the serine-and glycine-deficient diet, consumption of these amino acids must also be decreased to maintain steady-state circulating levels (Figure 4A). A number of circulating metabolites, including some amino acids, are cleared by TCA cycle oxidation driven by mass action^32^. We hypothesized that suppressed circulating levels of serine and glycine, induced by the dietary deficiency, leads to decreased catabolism by the SHMT2-serine dehydratase-TCA cycle pathway. To test this, we measured liver labeling of TCA cycle intermediates from U13C-glycine and U13C-serine infusions on control or ser/gly-free diets. The malate fractional labeling from both U13C-glycine and U13C-serine was decreased in mice on the deficient diet (Figure 5A). Consistent with mass action clearance, liver malate labeling from U13C-glycine has a linear relationship across physiological glycine concentrations (Figure 5B). This flux from glycine into the TCA cycle depends on SHMT catalytic activity, as it is ablated by SHIN2 treatment (Figure 5B). Thus, de novo synthesis from glucose is constant across circulating serine levels, and homeostasis instead requires variable glycine clearance by hepatic SHMT2 for TCA cycle oxidation (Figure 5C).

**Figure 5:**
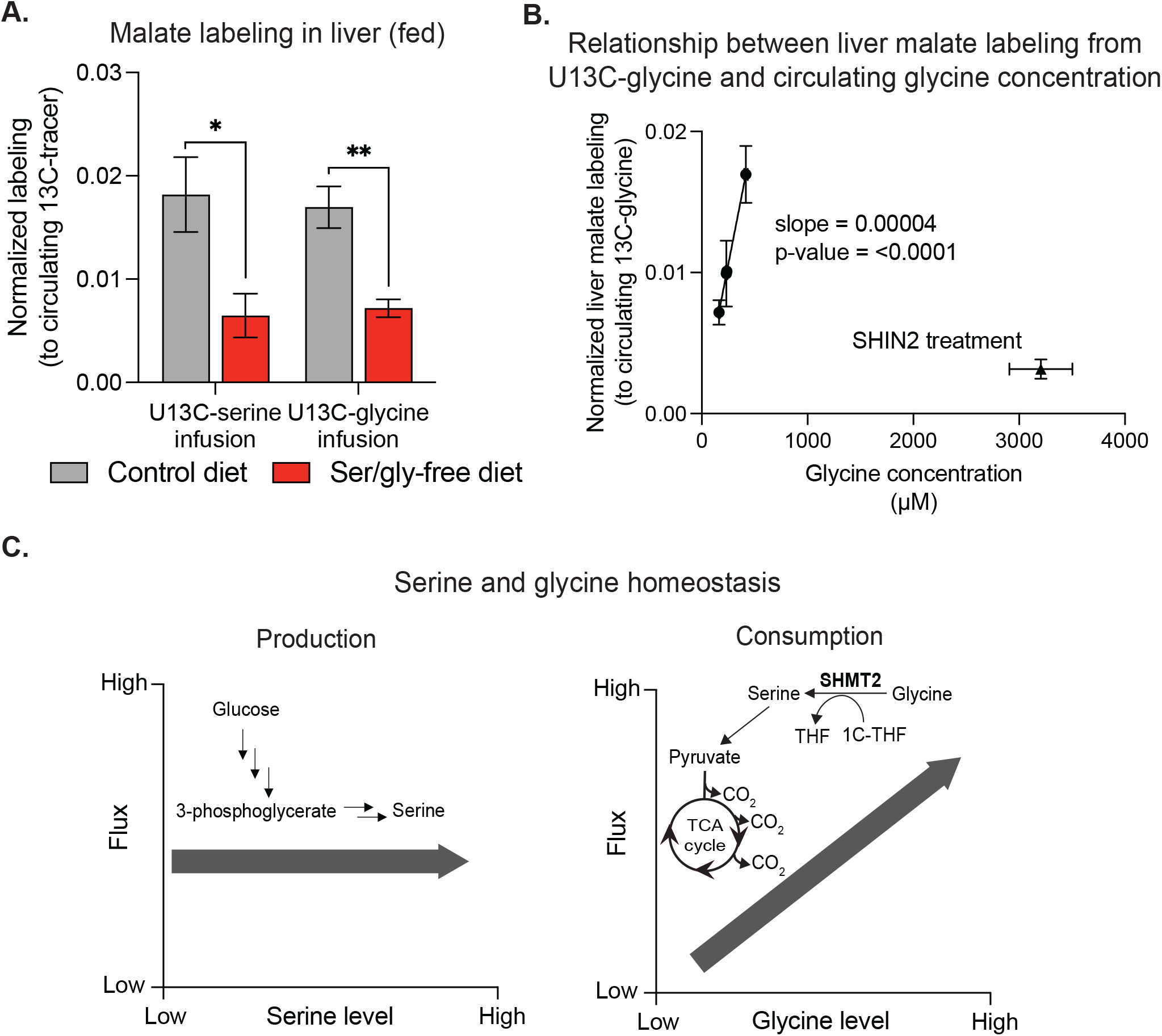
Serine and glycine homeostasis is achieved by mass-action driven TCA oxidation. (A) Normalized labeling from U13C-serine and U13C-glycine infusions of malate in liver for fed mice on control or ser/gly-free diet (n=4-6). (B) Relationship between U13C-glycine labeling of liver malate and circulating glycine concentration in mice across dietary conditions compared to mice treated with SHIN2. (C) Schematic of production from de novo serine synthesis (left) and SHMT2-mediated glycine consumption (right) pathways and the relationship of these pathway fluxes with varying serine and glycine levels. Proportionality between consumption flux and concentration drives homeostasis. All data are reported as mean ± SEM, p value by unpaired T test and replicates (n) indicated.

## DISCUSSION

This study finds that a primary route of mammalian glycine clearance is hepatic conversion to serine via reverse SHMT2 flux. This reverse flux through liver SHMT2 occurs in proportion to substrate levels, following mass action and thereby facilitating glycine homeostasis without substantial changes in biosynthetic flux.

All chemical reactions can occur in both the forward and reverse directions. The laws of thermodynamics dictate that net flux is in the direction with a negative Gibbs free energy change^33^. In metabolism, compartmentalization enables different reaction direction across organelles, cell types, or tissues. Glucose consumption by muscle glycolysis and production by hepatic gluconeogenesis is a classic example of opposite flux directions across tissues. Here we show the direction of SHMT flux is similarly tissue-dependent. Reverse SHMT2 flux in the liver occurs in concert with 1C-unit generating forward flux in proliferative cells like activated T cells. Such 1C-generating flux is of great physiological importance, but apparently lesser in absolute magnitude than SHMT2-mediated liver glycine consumption.

There is evidence of this net reverse SHMT1/2 flux in humans. Healthy and obese volunteers were infused with [U13C]glycine and [2,3,3-2H3]serine tracers to explore the impact of obesity on glycine metabolism^34^. While this work focused on flux differences between the two group and did not discuss the potential for whole-body SHMT flux to be net glycine consuming, the authors report quantitative fluxes from serine to glycine, and vice versa. Notably, the greater flux, by about two-fold (123 μmol/kg lean body mass/h), is in the glycine-consuming direction.

Both SHMT2 and GCS clear hepatic glycine within mitochondria. It is feasible that these reactions together form a single clearance pathway consuming two glycine molecules by using the 1C unit from GCS as a substrate for SHMT2^35^. We find, however, that GCS loss does not block hepatic serine labeling from U13C-glycine. Moreover, U13C-glycine infusions in humans show that GCS provides less than half of the 1C units consumed by reverse SHMT flux^34,36^. Other sources of mitochondrial 1C units in liver include choline and betaine.

In cancer cell lines, removal of serine and glycine from culture media increases de novo synthesis from glucose^5,11,37^. This occurs both by increased glycolytic carbon supply due to allosteric suppression of PKM2 activity and by transcriptional activation of serine synthesis pathway genes by activating transcription factor 4 (ATF4)^5,10,38^. In vivo, however, we observed no induction of de novo serine or glycine synthesis in response to removal of these nonessential amino acids from the diet. One interpretation is that serine and glycine levels do not fall low enough to lead to uncharged tRNA and ATF4 induction. In an effort to trigger such responses, we also examined mice on a low protein (5%) diet, where insufficiency of essential amino acids has been reported to activate, via uncharged tRNA, integrated stress response kinase GCN2 leading to ATF4 expression, but again we did not observe increased de novo serine or glycine synthesis^39^.

Consistent with endogenous serine and glycine metabolism remaining largely unaltered in response to diets lacking these amino acids, even the levels of serine and glycine stay the same except after eating, when they are lower due to lack of dietary influx. These observations are particularly important as serine and glycine-free diets have anticancer activity in mice and are currently being tested in humans (NCT05183295)^11,30,31,40–43^. Our data suggest that the impact of these diets, at least in mice, is limited to the fed state.

Given that production fluxes do not change, how is homeostasis maintained in response to serine and glycine-free diet? The simplest possibility is suppressed mass action-mediated consumption. Elevated levels of many metabolites lead, in linear proportion, to their enhanced consumption^32^. Here we find that suppressed serine and glycine levels, due to the dietary absence, lead in linear proportion to suppressed consumption. These data thus further support mass action as a central homeostatic mechanism in mammalian metabolism.

Proper glycine homeostasis is strongly associated with metabolic health. Humans with obesity, type II diabetes and non-alcoholic fatty liver disease have modestly decreased circulating glycine concentrations (down 10-20%)^44^. More generally, glycine level has an inverse association with BMI and weight loss interventions consistently increase glycine, tending to restore healthy levels^34,45,46^. Despite systematic studies in both rodents and patients, it remains unclear if impaired de novo synthesis or elevated catabolism drives this glycine depletion^47–49^. Studies on glycine catabolism in metabolic syndrome have focused on GCS flux. GCS flux was unchanged in fatty liver disease patients compared to healthy individuals^50^. Our work here uncovers glycine consumption by SHMT2. Investigation of whether this pathway is enhanced by metabolic syndrome merits exploration.

## ACKNOWLEDGEMENTS

M.J.M is funded by NIH grant F32CA250190. This work was funded by NIH Pioneer award DP1DK113643 and Ludwig Cancer Research to J.D.R. We are grateful to members of the Rabinowitz group for scientific discussions and insights.

## AUTHOR CONTRIBUTIONS

M.J.M and J.D.R. conceived and designed the study. M.J.M performed experiments and data analysis. C.H. performed catheterization surgeries and tail vein injections. M.J.M and J.D.R. wrote the manuscript.

## DECLARATION OF INTERESTS

J.D.R. is a member of the Rutgers Cancer Institute of New Jersey and the University of Pennsylvania Diabetes Research Center; a co-founder and stockholder in Empress Therapeutics and Serien Therapeutics; and an advisor and stockholder in Agios Pharmaceuticals, Bantam Pharmaceuticals, Colorado Research Partners, Rafael Pharmaceuticals, Barer Institute, and L.E.A.F. Pharmaceuticals.

## METHODS

### LEAD CONTACT AND MATERIALS AVAILABILITY

Further information and requests for resources and reagents should be directed to and will be fulfilled by the Lead Contact, Joshua D. Rabinowitz.

### EXPERIMENTAL MODEL AND SUBJECT DETAILS

Mouse and rat studies followed protocols approved by the Animal Care and Use Committee for Princeton University. C57BL/6N mice were obtained from Charles River Laboratories. *Shmt2^flox/flox^* mice in the C57BL/6 background were generated by Genome Editing Shared Resource at Rutgers University, and bred at Princeton University. *Shmt1^KO/KO^* mice were a generous gift from P.J. Stover and bred at Princeton University. *Gldc^flox/flox^* mice in the C57BL/6N background were obtained from Jackson Laboratory (strain #: 034928) and bred at Princeton University. Mice were allowed at least 5 days of acclimation to the facilities prior to experimentation, were randomly chosen for experimental groups and were on a normal light cycle (8 AM – 8 PM). Mouse experiments were performed on 10-14 week old animals. Animals received either a normal chow diet (PicoLab Rodent 20 5053 laboratory Diet St. Louis, MO) or an amino acid purified diet from TestDiet: Control (5WTV), serine and glycine-free (5B62), or low protein (5Z7D). Control and serine/glycine-free diets provide energy 15% from protein (as free amino acids), 17% from fat and 68% from carbohydrates. All amino acids were elevated proportionally to maintain total protein level in serine/glycine-free diet. Low protein diet provides energy 5% from protein (as free amino acids), 17% from fat and 78% from carbohydrates. Animals received purified diets 21 days prior to experimentation. For rat experiments, Sprague Dawley rats with arterial and jugular vein catheters were obtained at 5 weeks of age from Charles River Laboratory, received normal chow diet (PicoLab Rodent 20 5053 laboratory Diet St. Louis, MO) and acclimated to environment and human handling for 3 weeks prior to experiments.

## METHOD DETAILS

### Serum collection and tissue harvesting

Fasted samples were collected from mice 8 hours after food removal (chow removed at 9AM and sample collected at 5PM). Ad lib fed samples were collected at 8AM. Tail blood was collected in live mice by tail snip. Blood was directly collected into blood collection tubes with clotting factor (Sarstedt 16.442.100). Blood samples were stored on ice and then centrifuged at 16,000 × g for 10 min at 4°C to collect serum (top layer). Tissue harvest was performed after euthanasia by cervical dislocation. Tissues were quickly dissected in the order of liver, spleen, pancreas, kidney, small intestine, quadricep, diaphragm, heart, lung, brain and ears (skin). Tissues were clamped with a pre-cooled Wollenberger clamp in tin foil, and dropped in liquid nitrogen. Serum and tissue samples were stored at −80°C.

### Stable isotope infusions in mice

Aseptic surgery was performed to place a catheter in the right jugular vein and to connect the catheter to a vascular access button implanted under the skin on the back of the mouse. Catheters and vascular access buttons (VABs) were bought from Instech Labs. Mice were allowed to recover from jugular vein catheterization surgery for at least 5 days before experimentation. For intravenous infusions, 13C-metabolites (Cambridge Isotope Laboratories) were prepared in saline at following concentrations: 0.2 M (fasted) and 0.8 M (fed) U-13C-glucose (CLM-1396), 40 mM U-13C-glycine (CLM-1017), 30 mM U-13C-serine (CLM-1574), and 5 mM U-13C-valine (CLM-2249). The infusion setup (Instech Laboratories) included a swivel and tether to allow the mouse to move around the cage freely. Infusion rate was set to 0.1 μL/min/g body weight and tracer infused for 150 minutes followed by tail blood collection and tissue harvesting. Fasted infusions were collected at 5PM 8 hours after chow removal (started infusion at 2:30PM). Re-fed infusions were in mice fasted at 9AM, chow replaced and infusion started at 8PM, and infusion completed at 10:30PM.

### Stable isotope infusions in rats with arterial and venous blood collection

Chow was removed at 9AM and infusion finished at 5PM. 40 mM U-13C-glycine (CLM-1017) and 50 mM U-13C-serine (CLM-1574) was prepared in saline and infused at a rate of 0.1 μL/min/g body weight for 5 hours. Arterial blood was collected via catheter, and then the animal was anesthetized by pentobarbital administered through jugular vein catheter (55 mg/kg priming dose over one minute followed by 2.3 mg/kg/hr maintenance dose). After 20 minutes, rat was surgically opened and blood from renal, hepatic, portal and tail veins was collected using a syringe. Blood samples were placed on ice and centrifuged at 16,000 × g for 10 min at 4°C for serum collection.

### Pharmacological SHMT1/2 inhibition

The small molecule SHMT inhibitor SHIN2 was dissolved in 20% 2-hydroxypropyl-β-cyclodextrin as a 20 mg/ml stock and diluted in 0.9% saline to the desired concentration. Drug or vehicle control was intravenously infused through the jugular vein catheter at an infusion rate of 16.7 mg/kg/hr, which is equivalent to 200 mg/kg SHIN2 over a 12-hour period.

### Liver-specific gene knockout

*Shmt2^flox/flox^* mice or *Gldc^flox/flox^* mice along with wildtype litter mate controls were administered 2×10^11^ genomic copies of AAV8-*7BG*-Cre viral particles (Addgene, #107787) by tail vein injection (100 μL of 2×10^12^ genomic copies per mL stock). Experiments were carried out 21 days after virus injection.

### Metabolite extraction of serum

Serum (3 μl) was extracted with cold 100% methanol (40X), vortexed, and cooled on wet ice for ten minutes. For absolute quantification, a standard mix of N15-isotope labeled serine (51 μM) and glycine (206 μM) was spiked-in at equal volume to the serum sample (3 μl). Then, the extract was centrifuged at 16,000 × g for 10 minutes at 4°C and supernatant was transferred to tubes for LC-MS analysis.

### Metabolite extraction of tissues

Frozen tissue pieces were pulverized using a Cryomill (Retsch) at cryogenic temperature. Ground tissue was weighed (10–20 mg) and transferred into a precooled tube for extraction. Soluble metabolites extraction was done by adding −20 °C 40:40:20 methanol:acetonitrile:water with 1 mM N-ethylmaleimide (for alkylation of thiol containing metabolites) to the resulting powder (40 μl solvent per mg tissue) ^51^. For absolute quantification, a standard mix of isotope labeled metabolites of known concentration was spiked-in at a volume of 1 μl per mg tissue. Samples were vortexed for 10 seconds, cooled at 4°C (on wet ice) for 10 minutes and then centrifuged at 4 °C at 16,000 × *g* for 30 minutes. Supernatant was transferred to LC–MS vials for analysis.

### Metabolite measurement by LC-MS

LC–MS analysis for soluble metabolites was achieved on a quadrupole-orbitrap mass spectrometer (Thermo Scientific): the Q Exactive PLUS hybrid, Exploris 240 or Exploris 480. Each mass spectrometer was coupled to hydrophilic interaction chromatography (HILIC) via electrospray ionization. To perform the LC separation of serum and tissue samples, an XBridge BEH Amide column (150 mm × 2.1 mm, 2.5 μM particle size, Waters) was used with a gradient of solvent A (95%:5% H2O: acetonitrile with 20 mM ammonium acetate, 20 mM ammonium hydroxide, pH 9.4), and solvent B (100% acetonitrile). The gradient was 0 minutes, 85% B; 2 minutes, 85% B; 3 minutes, 80% B; 5 minutes, 80% B; 6 minutes, 75% B; 7 minutes, 75% B; 8 minutes, 70% B; 9 minutes, 70% B; 10 minutes, 50% B; 12 minutes, 50% B; 13 minutes, 25% B; 16 minutes, 25% B; 18 minutes, 0% B; 23 minutes, 0% B; 24 minutes, 85% B; 30 minutes, 85% B. The flow rate was 150 μl min^−1^, an injection volume of 10 μl for serum samples and 5 μl for tissue samples, and column temperature was 25°C. MS full scans were in negative or positive ion mode with a resolution of 140,000 at *m/z* 200 and scan range of 70–1,000 *m/z*. The automatic gain control (AGC) target was 1 × 10^6^. LC-MS peak files were analyzed and visualized with El-MAVEN (Elucidata) using 5 ppm ion extraction window, minimum peak intensity of 1 × 10^5^ ions, and minimum signal to background blank ratio of 2. For infusion experiments, the software package Accucor was used to correct for metabolite labeling from natural isotope abundance^52^.

### Definition of normalized metabolite labeling

When the 13C-labeled tracer X is infused, the normalized labeling of downstream metabolite Y is defined as *L_Y_/L_X_*, where *L_X_* and *L_Y_* are the fraction of labeled carbon atoms for metabolite X and Y defined as

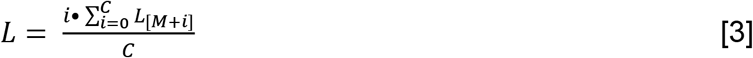

where *L_[M+i]_* is the fraction of mass isotopomer [M + i] and C is the total number of carbons in the metabolite. For infusions in mice, *L_X_* is the steady-state labeling in serum and *L_Y_* is the labeling in serum or tissues. For infusions in rats, *L_X_* is the labeling of blood into the liver (calculated based on 22% of arterial blood labeling and 78% portal vein blood labeling, both of which were directly measured) and *L_Y_* is the labeling of the blood draining from the liver (hepatic vein)^29^.

### Circulating turnover flux measurements

To measure the circulating (whole-body) turnover flux of a metabolite, we infused U13C-labeled form of the metabolite. At pseudo-steady state, we measured the mass isotope distribution of the metabolite in serum and the intact tracer circulatory turnover flux (*F_circ_*) was calculated as previously described. The fraction of the fully labeled tracer (i.e., the infused form), L_M+C_ (for example, glycine is M+2 due to having two carbon atoms) was used:

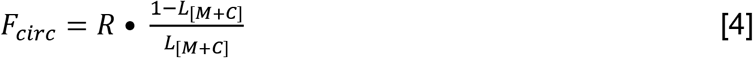

where R is the infusion rate of the labeled tracer. Since the turnover flux is a pseudo-steady state measurement, for minimally perturbative tracer infusions, production flux is approximately equal to consumption flux of the metabolite and thus *F_circ_* reflects both the circulating production and consumption fluxes of the infused metabolite.

### Contribution of non-essential amino acid production flux from protein degradation

Turnover flux of the essential amino acid valine was used to quantify production of the non-essential amino acids serine and glycine in the fasted state, as defined in equations [1] and [2]. Relative frequency (molar fraction) of valine (0.068), serine (0.081) and glycine (0.074) in protein was determined based on observed codon frequency in vertebrates^53^. Protein degradation is reduced in the fed state and amino acid production fluxes from protein degradation were taken to be 49% of fasted state fluxes based on prior literature data ^32^.

### Contribution of diet to amino acid production flux

Since amino acid purified chow have a defined quantity of serine and glycine, the proportion of turnover flux from diet in the fed state was calculated from quantity of chow consumed. The control chow (TestDiet, 5WTV) was 1.0% serine and 0.9% glycine by weight, so assuming 100% of amino acids consumed in chow reaches circulation, the production flux from diet (*F_AA←diet_*) is calculated by

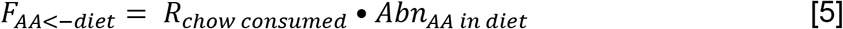

where *R_chow consumed_* is quantity of chow eaten by the mice per hour and *Abn_AA in diet_* is the molar abundance of the amino acid within the diet. All other production flux comes from endogenous sources, so endogenous production flux (*F_AA←endogenous_*) is calculated by

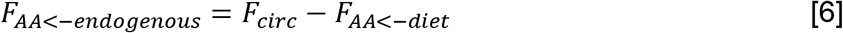

### De novo serine synthesis flux from circulating glucose

To measure the flux of serine carbons synthesized from circulating glucose, we performed U13C-glucose infusions and measured the normalized carbon labeling of serine in serum (*L_ser←glucose_*). From this, the flux of serine produced from glucose is calculated by

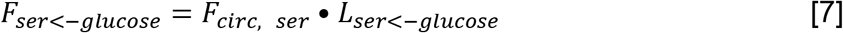

## QUANTIFICATION AND STATISTICAL ANALYSIS

Data are presented as means with error bars representing the standard error of the mean. A two-sided t-test was used to calculate p values and a p value of less than 0.05 was considered significant. For metabolomics, p value was corrected for multiple comparisons with a false discovery rate (FDR) cutoff of 0.1. Prism (GraphPad) was used for statistical analysis.

## SUPPLEMENTAL INFORMATION TITLES AND LEGENDS

**Figure S1:**
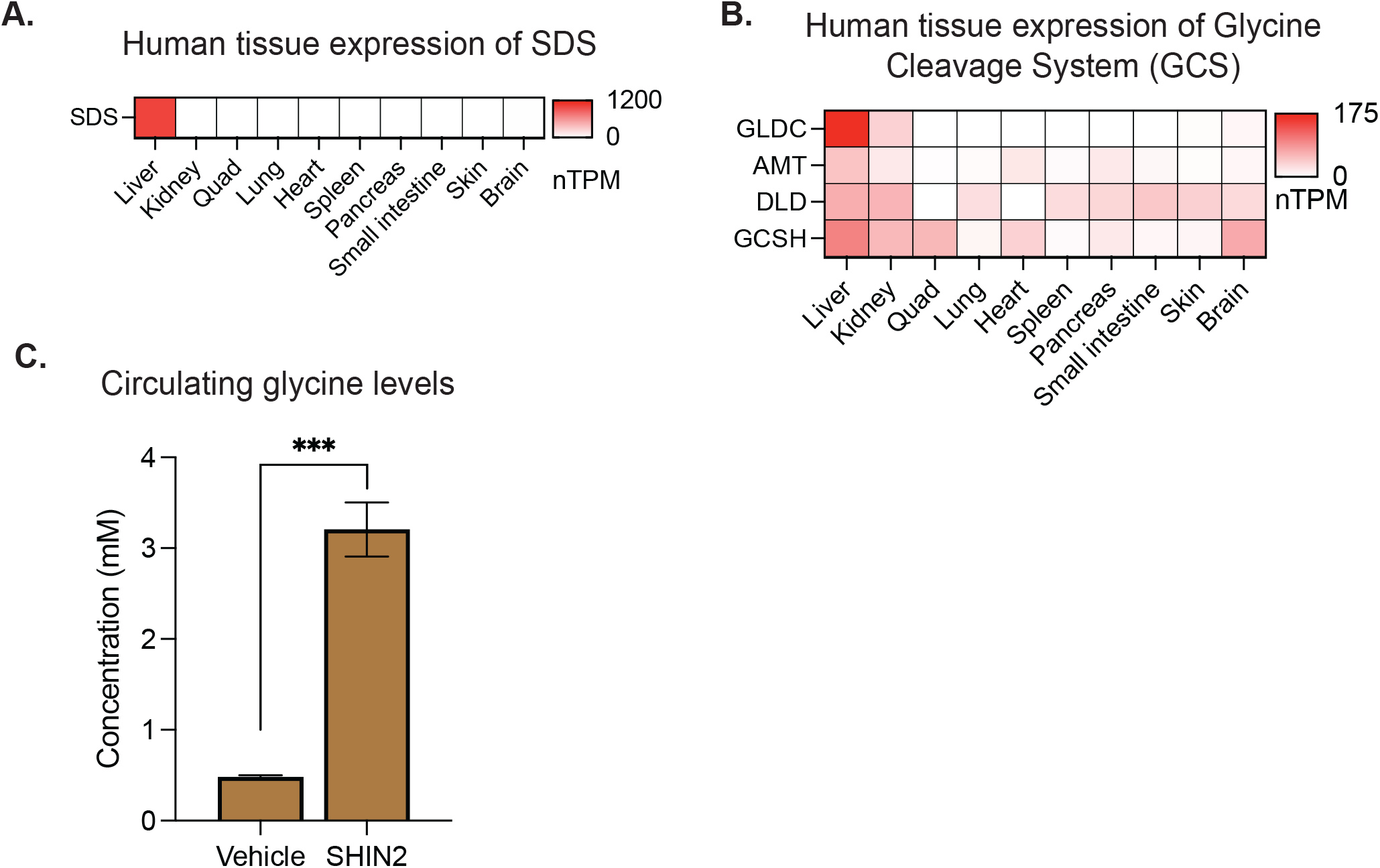
Liver-specific expression of serine and glycine catabolic enzymes. (A) Gene expression level of SDS across tissues in humans. (B) Gene expression level of enzymes that make up the glycine cleavage system (GCS) across tissues in humans. (C) Serum glycine concentrations from C57BL/6 mice treated with vehicle or SHIN2.

**Figure S2:**
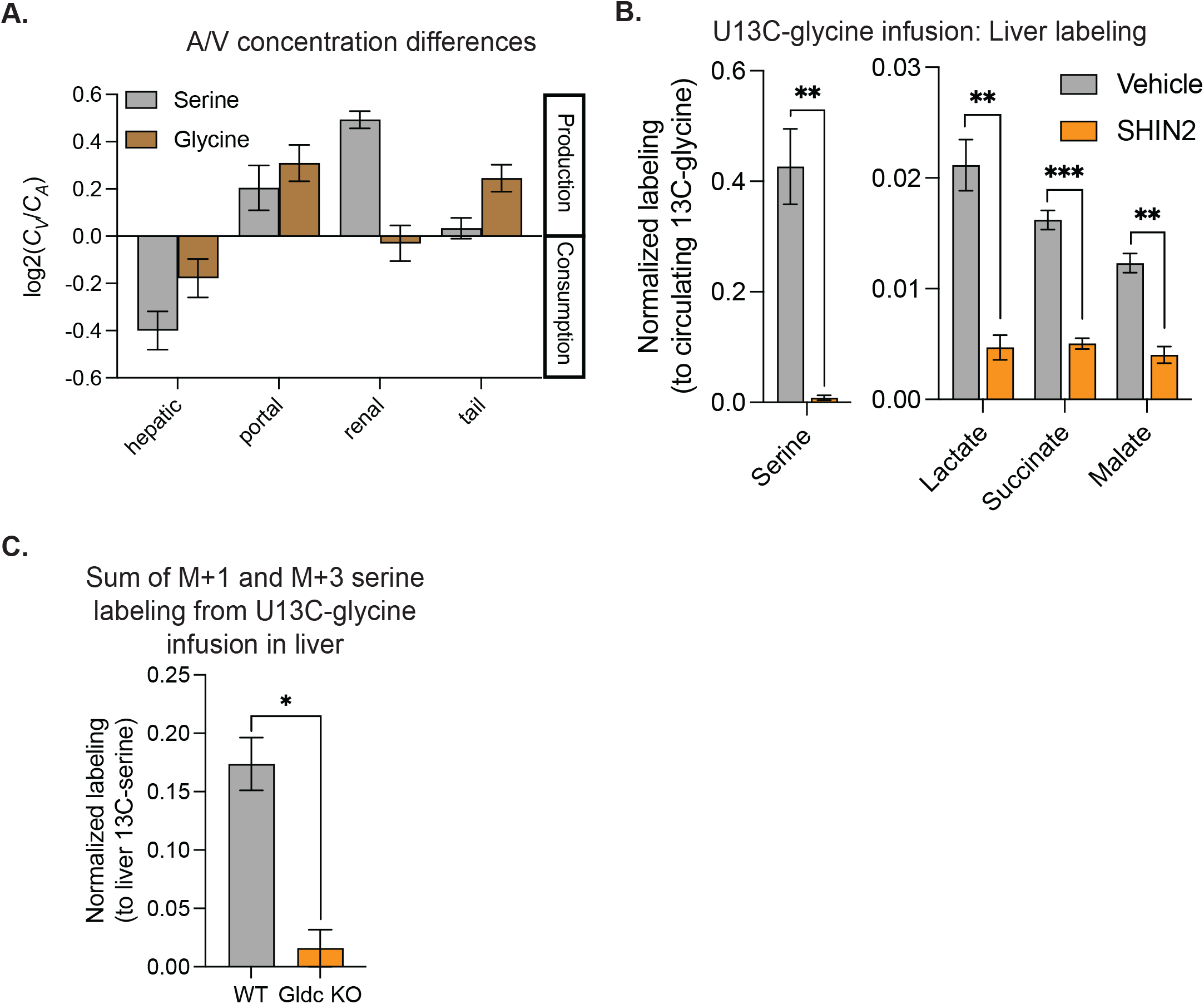
Hepatic glycine consumption is lost with SHMT1/2 inhibition. (A) Artery and venous concentrations of serine and glycine in Sprague Dawley rats (n=7-8). (B) Normalized labeling from U13C-glycine infusion of serine, lactate, succinate and malate in liver for fed mice treated with vehicle or SHIN2 (n=3). (C) Sum of M+1 and M+3 serine labeling from U13C-glycine infusion normalized to total liver serine labeling in wild-type and liver GLDC knockout mice.

**Figure S3:**
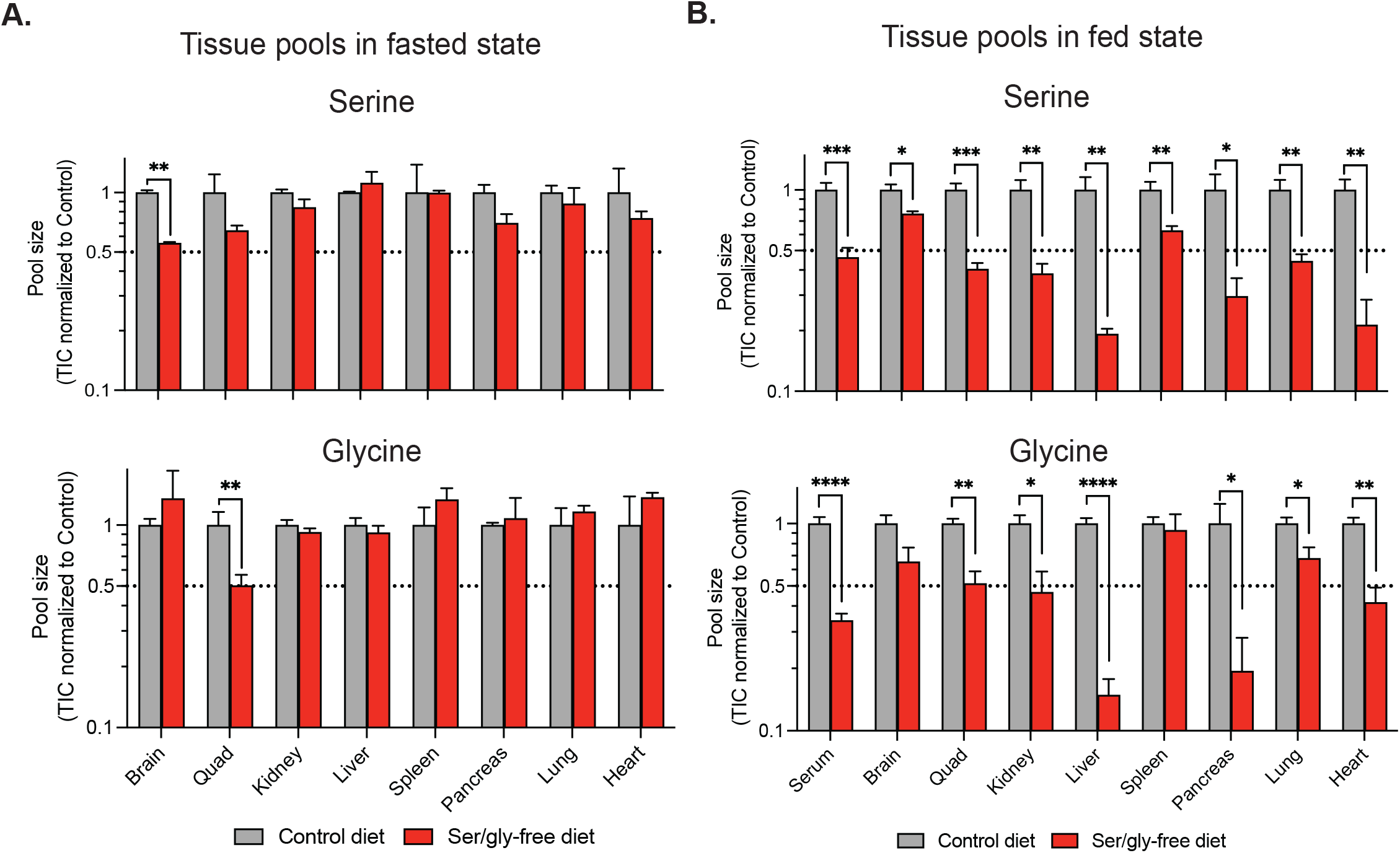
Serine and glycine tissue pools. (A) Fasted mice on control or ser/gly-free diet. (B) Fed.

**Figure S4:**
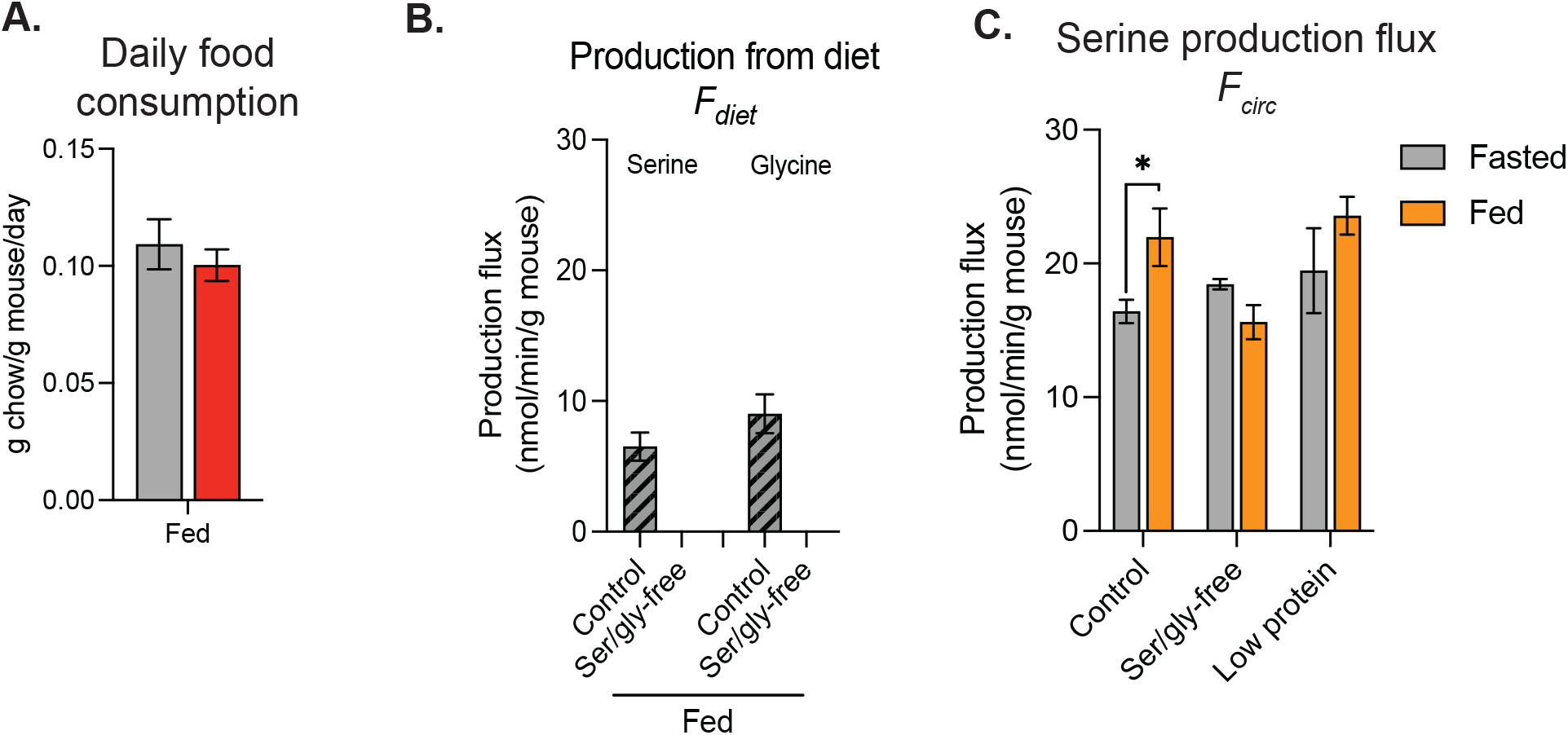
Quantifying amino acid production flux from diet. (A) Quantity of chow consumed by mice over 24 hours on control or ser/gly-free diet. (B) Based on quantity of chow consumed and proportion of chow that is serine or glycine, the production flux of each amino acid from diet in the fed state. (C) Serine turnover fluxes of fasted and fed mice on a control, ser/gly-free, and low protein diet.

